# Gene selection by incorporating genetic networks into case-control association studies

**DOI:** 10.1101/2022.03.10.483891

**Authors:** Xuewei Cao, Xiaoyu Liang, Shuanglin Zhang, Qiuying Sha

## Abstract

Large-scale genome-wide association studies (GWAS) have been successfully applied to a wide range of genetic variants underlying complex diseases. The network-based penalized regression approach has been developed to overcome the challenges caused by the computational efficiency for analyzing high-dimensional genomic data by incorporating a biological genetic network. In this paper, we propose a gene selection approach by incorporating genetic networks into case-control association studies for DNA sequence data or DNA methylation data. Instead of using traditional dimension reduction techniques such as principal component analyses and supervised principal component analyses, we use a linear combination of genotypes at SNPs or methylation values at CpG sites in each gene to capture gene-level signals. We develop three approaches for the linear combination: optimally weighted sum (OWS), LD-adjusted polygenic risk score (LD-PRS), and beta-based weighted sum (BWS). OWS and LD-PRS are supervised approaches that depend on the effect of each SNP or CpG site on the case-control status, while BWS can be extracted without using the case-control status. After using one of the linear combinations of genotypes or methylation values in each gene to capture gene-level signals, we regularize them to perform gene selection based on the biological network. Simulation studies show that the proposed approaches have higher true positive rates than using traditional dimension reduction techniques. We also apply our approaches to DNA methylation data and UK Biobank DNA sequence data for analyzing rheumatoid arthritis. The results show that the proposed methods can select potentially rheumatoid arthritis related genes that are missed by existing methods.

**Author Summary:** There is strong evidence showing that when genes are functionally related to each other in a genetic network, statistical methods utilizing prior biological network knowledge can outperform other methods that ignore genetic network structures. Therefore, statistical methods that can incorporate genetic network information into association analysis in human genetic association studies have been widely used since 2008. Here, we take advantage of recently developed methods to capture the gene-level signals in network-based penalized regression of high-dimensional genetic data. We have shown that the selection performance of our proposed methods can outperform three traditional principal component-based dimension reduction techniques in several simulation scenarios in terms of true positive rates. Meanwhile, by applying our methods in both DNA methylation data and DNA sequence data, the genes identified by our proposed methods can be significantly enriched into the rheumatoid arthritis pathway, such as genes *HLA-DMA*, *HLA-DPB1*, and *HLA-DQA2* in the HLA region.

## Introduction

With the maturation of modern molecular technologies, genomic data is increasingly available in large, diverse data sets (1). Those data sets give us an opportunity to use a large volume of human genetic data to explore meaningful insights about diseases. Over the last decade, large-scale genome-wide association studies (GWAS) have been successfully applied to a wide range of genetic variants underlying complex diseases (2). Different types of genetic variants have different biological functions in the human genome. Genotyping can identify small variations in DNA sequence within populations, such as single-nucleotide polymorphisms (SNPs) (3). Meanwhile, DNA methylation is an epigenetic maker that has suspected regulatory roles in a broad range of biological processes and diseases (4). Most penalized regression approaches have been developed to overcome the challenges caused by the computational efficiency for analyzing high-dimensional genomic data, such as elastic net (5), precision lasso (6), group lasso (7, 8), etc. However, Kim et al. (9) showed that these approaches ignore genetic network structures that have the worst selection performance in terms of the true positive rate.

The network-based regularization method has been shown that it can select phenotype related genes by incorporating genetic networks which can outperform other statistical methods that do not utilize genetic network information (9). Utilizing genetic network information indeed improves selection performance when genomic data are highly correlated among linked genes in the same biological process (i.e., genetic pathway). Therefore, the network-based regularization method has been developed in gene expression data (10) and DNA methylation data (11). To avoid the computational burden in analyzing high-dimensional genomic data, Kim et al. (9) proposed the approach that combines data dimension reduction techniques with network-based regularization to identify phenotype related genes. The dimension reduction techniques can capture the gene-level signals from multiple CpG sites or SNPs in a gene, such as the principal component (PC) based methods (PC, nPC, sPC, et al.) (9). PC method uses the first PC of DNA methylation data and nPC normalizes the first PC by the largest eigenvalue of the covariance matrix of methylation data. In addition, sPC uses the first PC of the data that only contains the CpG sites associated with the phenotype. It has been demonstrated that network-based penalized regression using PC-based dimension reduction techniques can outperform other methods that ignore genetic network structures (9) and the selection performance can be improved if the gene-level signals can capture more information.

Therefore, in this paper we propose a gene selection approach by incorporating genetic networks into case-control association studies with DNA sequence data or DNA methylation data. Instead of using traditional dimension reduction techniques such as PC-based methods, we use a linear combination of genotypes at SNPs or a linear combination of methylation values at CpG sites in each gene to capture gene-level signals. We develop three approaches for capturing gene-level signals: optimally weighted sum (OWS), LD-adjusted polygenic risk score (LD-PRS), and beta-based weighted sum (BWS). OWS and LD-PRS are supervised approaches that depend on the effect of each SNP or CpG site on the case-control status, while BWS can be extracted without using the case-control status. After we use one of the linear combinations of genotypes or methylation values in each gene to capture gene-level signals, we regularize them to perform gene selection based on the biological network. Simulation studies show that our proposed approaches have higher true positive rates than using traditional dimension reduction techniques. We also apply our approaches to DNA methylation data and UK Biobank DNA sequence data for rheumatoid arthritis patients and normal controls. The results show that the proposed methods can select potentially rheumatoid arthritis related genes that are missed by the PC-based dimension reduction techniques.

## Statistical Models and Methods

Consider a sample with *n* unrelated individuals, indexed by *i* = 1, 2,…, *n* . Support that there are a set of *M* genes in the analysis and a total of 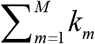 genetic components, such as SNPs in DNA sequence data or CpG sites in DNA methylation data, where *k_m_* is the number of genetic components in the *m^th^* gene. Let 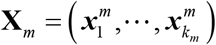 be an *n*×*k_m_* matrix of genetic components in the *m^th^* gene, where 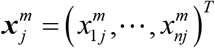 is the n-dimensional vector which represents the genetic data for the *j^th^* genetic component. Let ***y*** = (*y*_1_,…, *y_n_*)^*T*^ be an *n* ×1 vector of phenotype, where *y_i_* =1 denotes a case and *y_i_* = 0 denotes a control in a case-control study.

### Weighted linear combination methods

To capture gene-level signals from multiple genetic components in a gene, we develop three approaches using linear combinations of multiple genetic components in a gene: optimally weighted sum (OWS), LD-adjusted polygenic risk score (LD-PRS), and beta-based weighted sum (BWS). OWS and LD-PRS are the supervised methods based on the association between each genetic component and the phenotype, where OWS uses the optimal weight for each component (12), and LD-PRS can adjust for the LD structure of genetic components in a gene. BWS can be extracted without using the phenotype, where the genetic component is weighted by the beta distribution.

In OWS method, the OWS gene-level signal of the *m^th^* gene can be defined as

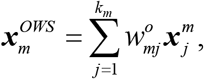

where 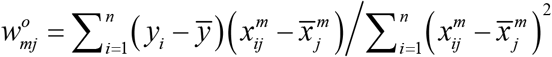 is the optimal weight for the *j^th^* genetic component in the *m^th^* gene; 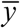 and 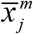 represent the sample mean of the phenotype and genetic data across all individuals, respectively. Large weight 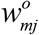 represents strong association between the genetic component and the phenotype (12).

To adjust for the influence of LD structure among the genetic components in a gene, the LD-adjusted genetic data can be defined as 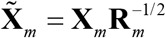, where **R**_*m*_ is the correlation matrix of **X**_*m*_ (13). Suppose 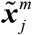 is the *j^th^* column of 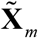, the LD-PRS gene-level signal of the *m^th^* gene is defined as

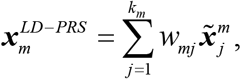

where 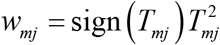 represents the weight of the *j^th^* genetic component in the *m^th^* gene and *T_mj_* is the score test statistic to test the association between the phenotype and the *j^th^* genetic component in the *m^th^* gene, which is given by

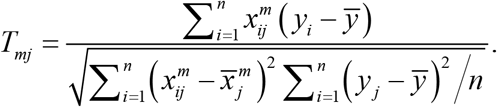

We define 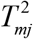 as the strength of the association and sign (*T_mj_*) as the direction of the effect. To make 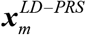 more robust, we use the following method to calculate 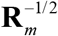. Let *λ*_1_ ≥ … ≥ *λ_k_m__* and ***e***_1_,…, ***e****_k_m__* be the eigenvalues and corresponding eigenvectors of the correlation matrix **R**_*m*_. Thus, the eigenvectors ***e***_1_,…, ***e****_k_m__* represent new orthogonal axes corresponding to decreasing variability of the genetic data. We only use the first *J* components to calculate 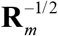 since very small values of eigenvalues can make 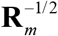 unstable, where *J* is the smallest number such that 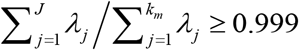. Therefore, 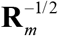 can be written as 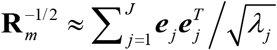.

In BWS method, the genetic component is weighted by the beta distribution which is similar to SNP-set (Sequence) Kernel Association Test (SKAT) (14). Decreasing the weight of noncausal genetic components and increasing the weight of causal components can yield improved power. The BWS gene-level signal of the *m^th^* gene is defined as

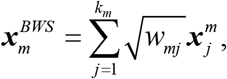

where 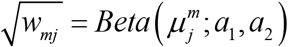 represents the weight of the *j^th^* genetic component in the *m^th^* gene which is extracted without using the phenotype. For DNA sequence data, 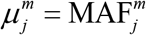 and the suggested settings of two parameters are *a*_1_ =1 and *a*2 = 25 because it increases the weight of rare variants (14), where 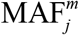 denotes the minor allele frequency of the *j^th^* genetic component in the *m^th^* gene. For DNA methylation data, 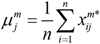 and 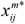 is the methylation *β* value for the *j^th^* CpG site of the *i^th^* individual and *a*_1_ = *a*_2_ = 0.5 corresponds to 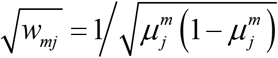.

Using one of the above weighted linear combination methods for each gene, the dimension of genetic components 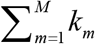 are reduced down to *M* gene-level signals. We use***z***_*i*_ = (*z*_*i*1_,…, *z*_*iM*_)^*T*^ to denote a gene-level signal as the feature of the *i^th^* individual across all genes, which can be obtained from one of the three methods, 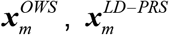, and 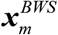.

### Network-based regularization

Consider **A** = (*a_mk_*) is an *M* × *M* adjacency matrix which represents the undirected network connections among genes, where *a_mk_* =1 represents the *m^th^* and *k^th^* genes are within the same biological set (i.e., pathway, etc.) and *a_mk_* = 0 otherwise. Let **D** = diag (*d*_1_,…, *d_M_*) be an *M* dimensional degree matrix, where the *m^th^* diagonal element is 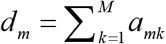 which represents the total number of genetic links of the *m^th^* gene. Therefore, the symmetric normalized Laplacian matrix **L** = **I** − **D**^-1/2^**AD**^-1/2^ represents a genetic network structure, where the elements of **L** are given by

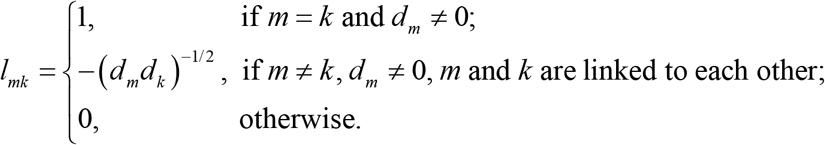

Let *β*_0_ and ***β*** = (*β*_1_,…,*β* _*M*_)^*T*^ be the intercept and the effect vector of *M* genes, respectively. The likelihood function of the phenotype is given by

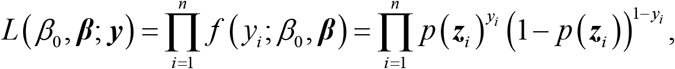

where *p* (***z***_*i*_) = Pr (*y_i_* = 1| ***z***_*i*_) represents the probability that the *i^th^* individual is a case, which can be calculated by

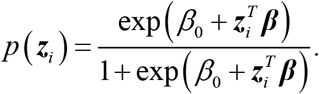

Based on the genetic network structure, the penalized logistic likelihood using network-based regularization (9) is given by

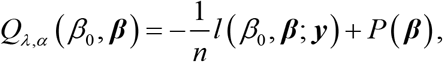

where *l* (*β*_0_, ***β***; ***y***) = log *L*(*β*_0_, ***β***; ***y***) is the log-likelihood function and *P* (***β***) is a penalty term which is a combination of the *l*_1_ penalty and squared *l*_2_ penalty incorporating the genetic network structure. *P* (***β***) is defined as

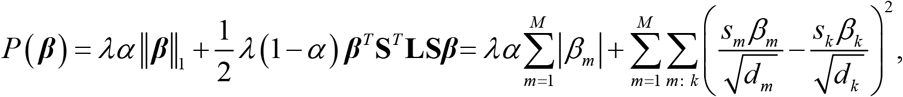

where ∥g∥_1_ is a *l*_1_ norm, and **S** = diag (*s*_1_,…, *s_M_*) is a diagonal matrix of the estimated signs of the regression coefficients on the diagonal entries *s_m_* ∈ {−1,1} for *m* = 1,…, *M*, which can be obtained from ordinary regression for *M* < *n*, and ridge regression for *M* ≥ *n* . If the genes have different effects on a phenotype, then the signs of the corresponding regression coefficients will be different, where the matrix **S** has been shown that it can accommodate the problem of failure of local smoothness between linked genes (15). *λ* is a tuning parameter that controls sparsity of the network-based regularization, *α*∈[0,1] is a mixing proportion between lasso penalty and network-based penalty, and *m* : *k* denotes that the *m^th^* and *k^th^* genes are linked to each other in the genetic network.

For a given pair of *λ* and *α*, we can estimate the interpret, *β*_0_, and the effect vector of *M* genes, ***β***, by minimizing the penalized logistic likelihood *Q_λ,α_*(*β*_0_, ***β***). It is not difficult to show the penalty function *P* (***β***) is convex (9, 16), so the solution *β*_0_ and ***β*** can be obtained via one of the convex optimization algorithms. We use the R package “pclogit” to estimate *β*_0_ and ***β*** which implements the cyclic coordinate descent algorithm (11, 17).

### Stability selection

The most common method to obtain the optimal tuning parameters *λ* and *α* is to use k-fold cross-validation. However, the results of selection are not stable due to that the samples are randomly split in the cross-validation (9). Meinshausen et al. (18) proposed a stability selection method that used a half-sample	approach in combination with selection algorithms.

In this paper, the half-sample method is used to compute the selection probability (SP) for each gene. Let *S_b_* be the *b^th^* random subsample that has a size of ⌊*n*/2⌋ without replacement, where ⌊g⌋ is a floor function which represents the greatest integer less than or equal to the value in the function. In each subsample set *S_b_*, there are ⌊*n*_cases_/2⌋ randomly subsampled cases and ⌊*n*_controls_/2⌋ controls, where *n*_cases_ and *n*_controls_ are the number of cases and controls in the data, respectively. For fixed values of *λ* and *α*, we estimate regression coefficient 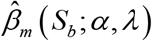 for the *m^th^* gene according to the above network-based regularization method based on the subsample set *S_b_*. Then, we repeat the half-sample method *B* times and count the total number of 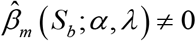 for *b* =1,…, *B*. The SP of the *m^th^* gene can be obtained based on grid sets of *α* and *λ*, which is computed by

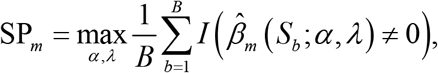

where 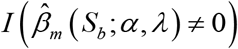 is an indicator function; 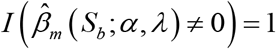 if 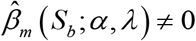 for *b* =1,…, *B*. The SP_*m*_ indicates that the maximum value of the proportion of the *m^th^* gene which has been selected using half-sample method *B* times among all choices of the tuning parameters *λ* and *α*. As suggested in R package “glmnet”, we consider a total of 600 pairs of tuning parameters *λ* and *α* to compute SP for each gene in simulation studies and real data analysis, where we choose six *α*values *α*= (0.1, 0.2,…, 0.6) and 100 *λ* values for each *α*. Let 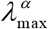 be the smallest value of *λ* for a given *α* such that all the coefficients are zeros (19). The grid set of *λ* for each *α*can be set as a log_10_-space from 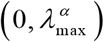.

There are two main advantages of the stability selection method. First, we do not need to select the optimal tuning parameters *λ* and *α*. After setting the two grid sets of *α*∈[0,1] and *λ*> 0, we estimate regression coefficient 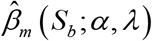 for the *m^th^* gene according to the above network-based regularization method for each pair of *λ* and *α*. Then, the SP is the largest proportion across all *λ* and *α*. Second, the half-sample method is used to obtain a stable selection result compared with the cross-validation. We use *B* = 100 in simulation studies and *B* = 500 in real data analysis.

### Simulation Studies

To evaluate if our proposed three weighted linear combination methods, OWS, LD-PRS, and BWS, outperform PC-based dimension reduction techniques, we follow the simulation settings in Kim et al. (9). The individual-level genetic data where linked genes within a biological network are correlated with each other are generated using the following three steps:

*Step 1: Construct an M dimensional covariance matrix from an arbitrary graph based on a Gaussian graphical model.*

Consider a total number of *M* = 1000 genes that contain 10 disjointed network modules, each of which consists of 100 genes. We construct each network module from Figure S1, which contains a centered gene correlated with other genes with a few links in one network module. Therefore, the adjacency matrix **A** = (*a_mk_*) of those 1000 genes is constructed based on the connections among genes in each network module, where *a_mk_* =1 represents the *m^th^* and *k^th^* genes are within the same network module and *a_mk_* = 0 otherwise. Next, we apply a Gaussian graphical model to generate a covariance matrix of 1000 genes (20). Following the settings in Peng et al. (20), the initial concentration matrix **Ω** = (*ρ_mk_*)_*M*×*M*_ is generated by

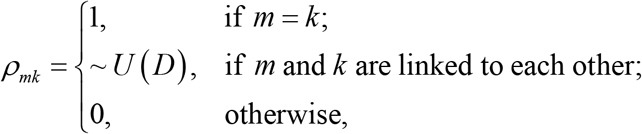

where *D* = [−0.7, −0.1]∪[0.1, 0.7] and *U* (*D*) represents a random variable from a uniform distribution on the domain *D* . We then rescale the non-zero elements in the concentration matrix to assure positive definiteness, that is, we divide each off-diagonal element by 1.5-fold of the sum. Finally, we average the rescaled matrix with its transpose to ensure symmetry and set the diagonal entries to be one. We denote the final matrix as **Ω** and the covariance matrix **Σ** can be determined by 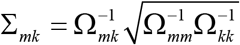, where 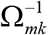 represents the (*m*, *k*)^*th*^ element of the inversed concentration matrix **Ω**^−1^. Note that the correlations between linked genes are much higher than that of unlinked genes.

*Step 2: Generate M gene-level signals from different multivariate normal distributions for cases and controls, respectively.*

In this step, we consider two scenarios to set up the phenotype-related genes. In scenario 1, we assume that only 45 genes in the first network module are phenotype related, where these 45 genes contain the centered gene and four subgroups of genes denoted as *g*_1_, *g*_2_, *g*_3_, and *g*_4_, respectively. In scenario 2, we assume that 48 genes in the first four network modules are phenotype related, where each of network modules contains one centered gene and a subgroup of genes which are denoted as *g*_1_, *g*_2_, *g*_3_, and *g*_4_, respectively. For each scenario, let ***μ*** = (*μ*_1_,…,*μ_M_*)^*T*^ be the mean vector, where ***μ*** = (0,…, 0)^*T*^ for the control group. In the case group, we set *μ_m_* = 0 for neutral genes (i.e., 955 genes in scenario 1 and 952 genes in scenario 2). In contrast, the mean of phenotype related genes *μ_m_* is defined as

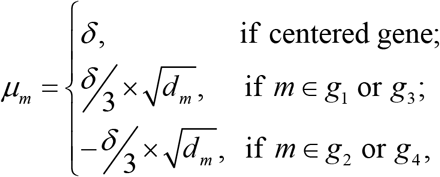

where *δ* is the strength of association signals and *d_m_* is the total number of genetic links for the *m^th^* gene. Therefore, the gene-level signals for each individual can be generated from a multivariate normal distribution, ***z***_*i*_ : *MVN*(***μ***, **Σ**) for *i* = 1,…, *n*.

*Step 3: Generate DNA methylation and DNA sequence data based on each gene-level signal.*

Consider *k_m_* = 10 genetic components for *m* = 1,…, *M* and a total of 10,000 genetic components in simulation studies. In this step, we consider two types of genetic data, DNA methylation data and DNA sequence data. Let *ω* be the number of components correlated with the gene-level signal value *z_im_* for the *i^th^* individual and the *m^th^* gene, which controls the number of causal or neutral components. The methylation value of the *i^th^* individual and *j^th^* CpG site in the *m^th^* gene is denoted by 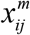 which can be generated by

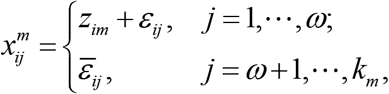

where *ε_ij_* : *N* (0,*σ*^2^) indicates the difference between the *j^th^* CpG site and gene-level signal *z_im_* and *σ*^2^ is the error variance that controls the noise level of association signals. 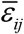 follows a normal distribution with mean 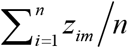 and variance *σ*^2^.

The value of genotype data usually indicates the genotypic score of an individual at a SNP which is the number of minor alleles that the individual carries at a SNP. We first generate two different continuous data 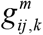 for *k* = 1 or 2 to indicate the genotypic value for two alleles which are defined as

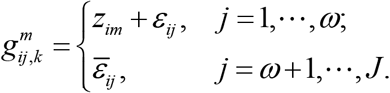

Next, we convert continuous data 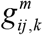 to binary data 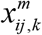 based on a fixed 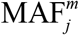 for the *j^th^* SNP in the *m^th^* gene. Finally, the genotype data 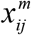 for the *j^th^* SNP in the *m^th^* gene is generated by 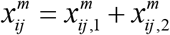. Therefore, the genotype data is coded as 0, 1, or 2. In our simulation studies, we consider five rare variants and five common variants in each gene, where the MAFs for rare variants are randomly generated from a uniform distribution *U* (0.001, 0.01) and MAFs for common variants are from *U* (0.01, 0.5) .

After generating the individual-level DNA methylation data and DNA sequence data based on a biological network structure from the above three steps, we use the proposed three weighted linear combination methods, OWS, LD-PRS, and BWS, to calculate the gene-level signals ***z*** = (*z*_*i*1_,…, *z_iM_*)^*T*^ for the *i^th^* individual across all genes. Then, the selection probability for each gene can be obtained by using a half-sample method 100 times and the network-based regularization method across 600 pairs of tuning parameters *λ* and *α*. The true positive rate (TPR) is used to evaluate the selection performance, which is defined as the number of selected top-ranked genes among the phenotype related genes divided by the total number of phenotype related genes. For each scenario, we consider a total of *n* = 1000 individuals which contain 500 cases and 500 controls for the balance case-control studies. Figs 1-2 show the TPR comparisons for the balance case-control studies in scenario 1. We compare our proposed three linear combination approaches with three PC-based dimension reduction techniques: PC, nPC, and sPC, which have been shown higher TPR than other methods that do not utilize biological network information. We first compute selection probabilities of all genes and then rank top genes based on the selection probabilities for each method.

**Fig 1.**
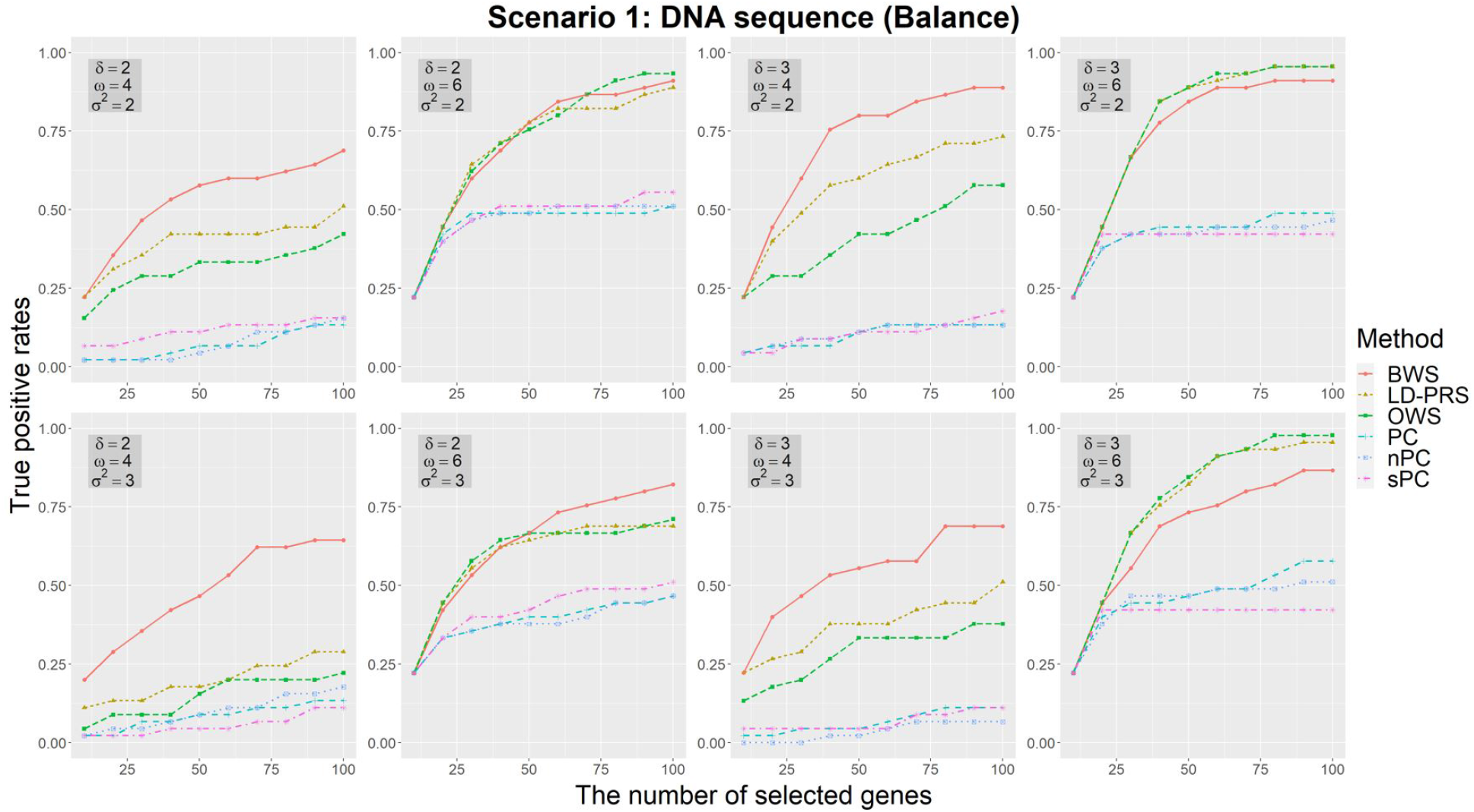
The true positive rates of the methods based on different gene-level signals for balance case-control studies with DNA sequence data in scenario 1, where there are five rare variants and five common variants in each gene. According to the different number of selected top genes, three parameters are used to vary the genetic effect: the strength of association signals *δ*, the number of SNPs in each gene related to gene-level signals *ω*, and the noise level of association signals *σ*^2^. The selection probabilities are calculated using half-sample method 100 times.

**Fig 2.**
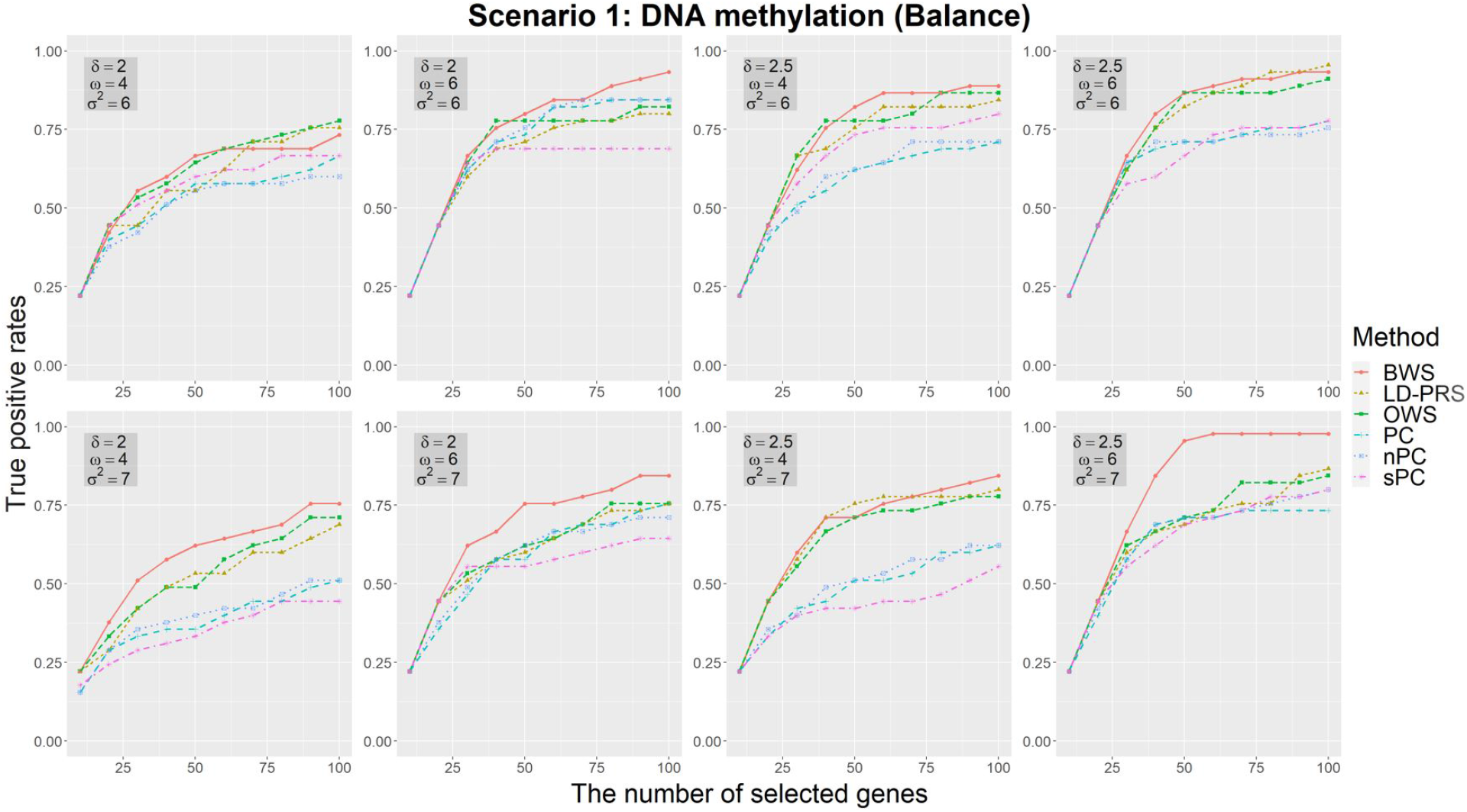
The true positive rates of the methods based on different gene-level signals for balance case-control studies with DNA methylation data in scenario 1. According to the different number of selected top-genes, three parameters are used to vary the genetic effect: the strength of association signals *δ*, the number of CpG sites in each gene related to gene-level signals *ω*, and the noise level of association signals *σ*^2^. The selection probabilities are calculated using half-sample method 100 times.

In DNA sequence data analysis (Fig 1), we pre-set the strength of association signals (*δ*= 2, 3), the number of components correlated with the gene-level signal (*ω*= 4, 6), and the error variance which controls the noise level of association signals (*σ*^2^ = 2,3). The proposed OWS, LD-PRS, and BWS have the best selection performance in all eight simulation settings according to TPR. When the number of causal SNPs in a gene is small (*ω*= 4), BWS has the uniformly highest TPR regardless of the size of the error variance. However, TPRs of the supervised approaches, OWS and LD-PRS, are higher than or similar as that of the unsupervised approach, BWS, when the number of SNPs in a gene is large (*ω*= 6). Overall, BWS shows the best selection performance in all simulation settings for DNA sequence data analysis. LD-PRS is better than OWS due to LD-PRS can adjust for the LD structure of the SNPs. In DNA methylation data analysis (Fig 2), we pre-set *δ*= 2, 2.5, *ω*= 4, 6, and *σ*^2^ = 6,7 . All methods have similar performance according to TPR when the strength of the association signal is small (*δ*= 2). However, the proposed three methods have higher TPRs than PC-based methods when the strength of the association signal is large (*δ*= 2.5). Particularly, when the number of components correlated with the gene-level signal is large (*ω*= 6), BWS has the uniformly highest TPR regardless of the size of the error variance and the strength of association signals. BWS also shows the best selection performance in all simulation settings for DNA methylation data analysis. LD-PRS and OWS have similar performance but have higher TPRs than the other three PC-based methods.

Figs S2-S3 show the TPR comparisons for the balance case-control studies under scenario 2. The patterns of TPR comparisons under scenario 2 for DNA methylation data and DNA sequence data are similar to those under scenario 1 (Figs 1-2). Meanwhile, we also perform TPR comparisons for the unbalance case-control studies, where there are a total of *n* = 1000 individuals with 100 cases and 900 controls. Figs S4-S7 show the TPR comparisons for the unbalance case-control studies. The patterns of TPR comparisons under these two scenarios for DNA methylation data and DNA sequence data are similar to those observed in Figs 1-2 and Figs S2-S3.

### Applications

To evaluate the performance of our proposed three approaches for the linear combination in real data analyses, we apply our approaches to DNA methylation data and UK Biobank data for DNA sequence of rheumatoid arthritis (RA) patients and normal controls. In order to utilize biological network information, we employ the same pathway information as in Kim et al.(9), where there are seven genetic network databases from Biocarta, HumnaCyc, KEGG, NCI, Panther, Reactome, and SPIKE in R package ‘graphite’. There are 11,381 linked genes in the package, of which 672,571 edges among those genes are in the biological network.

Due to the outperformance of the nPC (9) compared with the other PC-based dimension reduction techniques, PC and sPC, we only apply nPC to compare the performance with our proposed methods in real data analyses. We consider all genes according to the USCS (GRCh37/hg19) genome sequence annotation list which can be downloaded from the UCSC website (https://hgdownload.soe.ucsc.edu/goldenPath/hg19/bigZips/). There are 28,152 genes with gene symbols from the UCSC database.

### Application to DNA methylation data

The DNA methylation data was measured using the Illumina HumanMethylation450 BeadChip from 354 RA patients (cases) and 335 normal controls (21, 22). The dataset can be found in the NCBI Gene Expression Omnibus (GEO; https://www.ncbi.nlm.nih.gov/geo/) with identifier GSE42861, where the methylation *β* values of CpG sites are provided. Then, we convert *β* values to *M* values using logit (base 2) function for further analysis. We find that only 10,737 linked genes matched with genes in the above biological network that contain at least one CpG site. After pruning CpG sites in each gene, we calculate gene-level signals using OWS, LD-PRS, BWS, and nPC. Then, the selection probability for each gene can be obtained by using half-sample method 500 times and the network-based regularization method across 600 pairs of tuning parameters *λ* and *α*.

We select the top 100 genes according to the selection probabilities of each method. We search the GWAS catalog (https://www.ebi.ac.uk/gwas/) for genes that are associated with RA. Table 1 shows the genes in the GWAS catalog that are also identified by OWS, LD-PRS, BWS, and nPC. OWS identifies 11 genes, LD-PRS identifies 12 genes, BWS identifies 8 genes, and nPC identifies 10 genes. Meanwhile, the number of overlapped genes by each method in the DNA methylation data analysis is summarized in Fig S8. There are four genes identified by all of these four methods, *HLA-DQA2*, *HLA-DRB1*, *HLA-DQB1*, and *CD1C*. Gene *HLA-DRB1* (23-29) and gene *HLA-DQB1* (23, 29-35) play a central role in the immune system and have been reported in the GWAS catalog. No literature reported gene *HLA-DQA2* that was significantly associated with RA in GWAS catalog. However, the SPs of gene *HLA-DQA2* calculated by our proposed three methods, OWS, LD-PRS, and BWS, are all 1.000. Also, the SP of gene *HLA-DQA2* is 0.852, which is also on the top 100 genes identified by nPC method. Notably, gene *HLA-DQA2* is in the rheumatoid arthritis pathway (KEGG: hsa05323) and the literature (36) has shown that genes in the human leukocyte antigen (HLA) region remain the most powerful disease risk genes in RA.

**Table 1.**
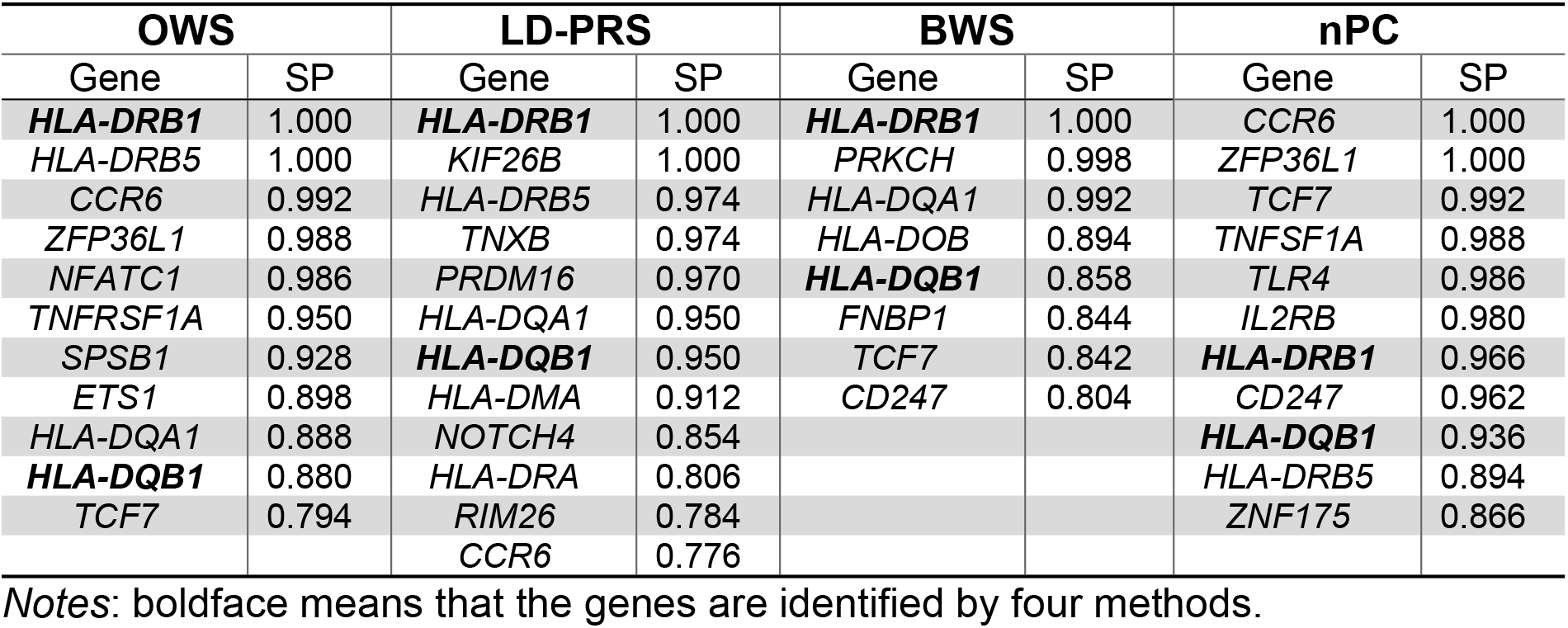
GWAS catalog reported genes identified by OWS, LD-PRS, and BWS in DNA methylation data.

In order to better understand the biological meaning behind the top 100 selected genes by each method, we map those genes to the Kyoto Encyclopedia of Genes and Genomes (KEGG) pathways using a functional annotation tool named Database for Annotation, Visualization, and Integrated Discovery Bioinformatics Resource (37, 38) (DAVID: https://david.ncifcrf.gov/) for pathway enrichment analysis. In this study, significantly enriched pathways are identified by the top 100 selected genes if FDR < 0.05. In Fig 3, there are 21 significantly enriched pathways identified by OWS, BWS, and LD-PRS, in which the RA pathway is significantly enriched with FDR_OWS_= 1.48E-04, FDR_BWS_= 7.80E-03, and FDR_LD-PRS_= 8.03E-07, respectively; RA pathway is also significantly enriched in a total of 18 pathways identified by nPC with FDR_nPC_= 2.91E-03.

**Fig 3.**
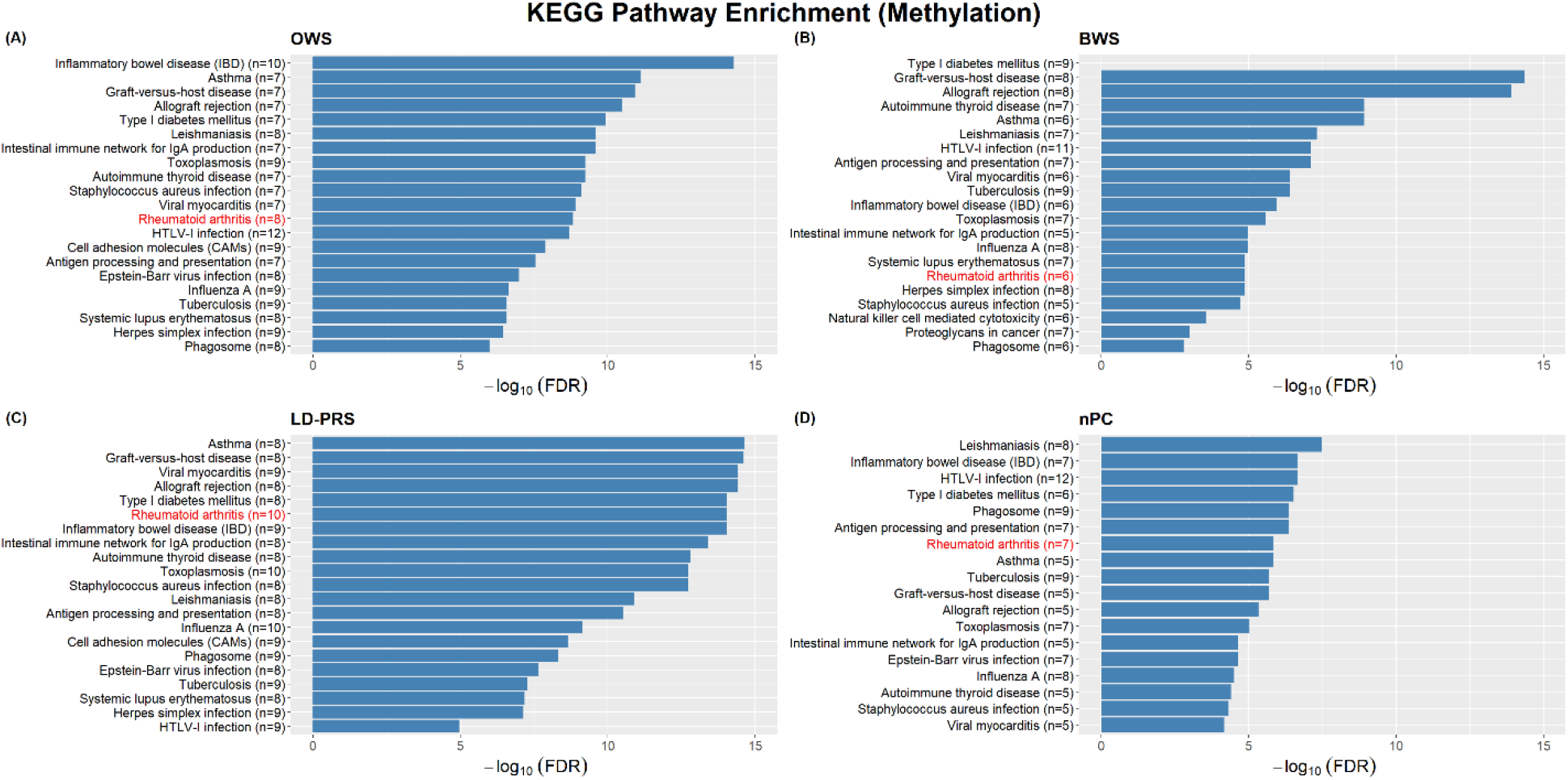
The KEGG pathway enrichment analysis results of BWS, LD-PRS, OWS, and nPC for DNA methylation data.

The overlapping genes between the top 100 genes identified by each method and genes in RA pathway are shown in Fig 4. The number below each method indicates the total number of overlapping genes identified by the corresponding method and genes in RA pathway. LD-PRS has the smallest pathway enriched FDR and identifies the most overlapping genes (n=10); genes *HLA-DMA* (SP = 0.912) and *LTB* (SP = 0.998) are uniquely identified. OWS identifies eight overlapping genes which contain one unique gene *HLA-DPB1* (SP = 0.85); meanwhile, BWS identifies six overlapping genes that contain two unique genes *TNF* (SP = 0.980) and *HLA-DOB* (SP = 0.894). Comparing the results of our proposed three methods with nPC, five HLA-family genes (*HLA-DMA*, *HLA-DOB*, *HLA-DPB1*, *HLA-DPA1*, and *HLA-DQA1*) and two RA pathway genes (*LTB* and *TNF*) are uniquely identified. The results show that the proposed methods can select potentially RA related genes that are missed by nPC.

**Fig 4.**
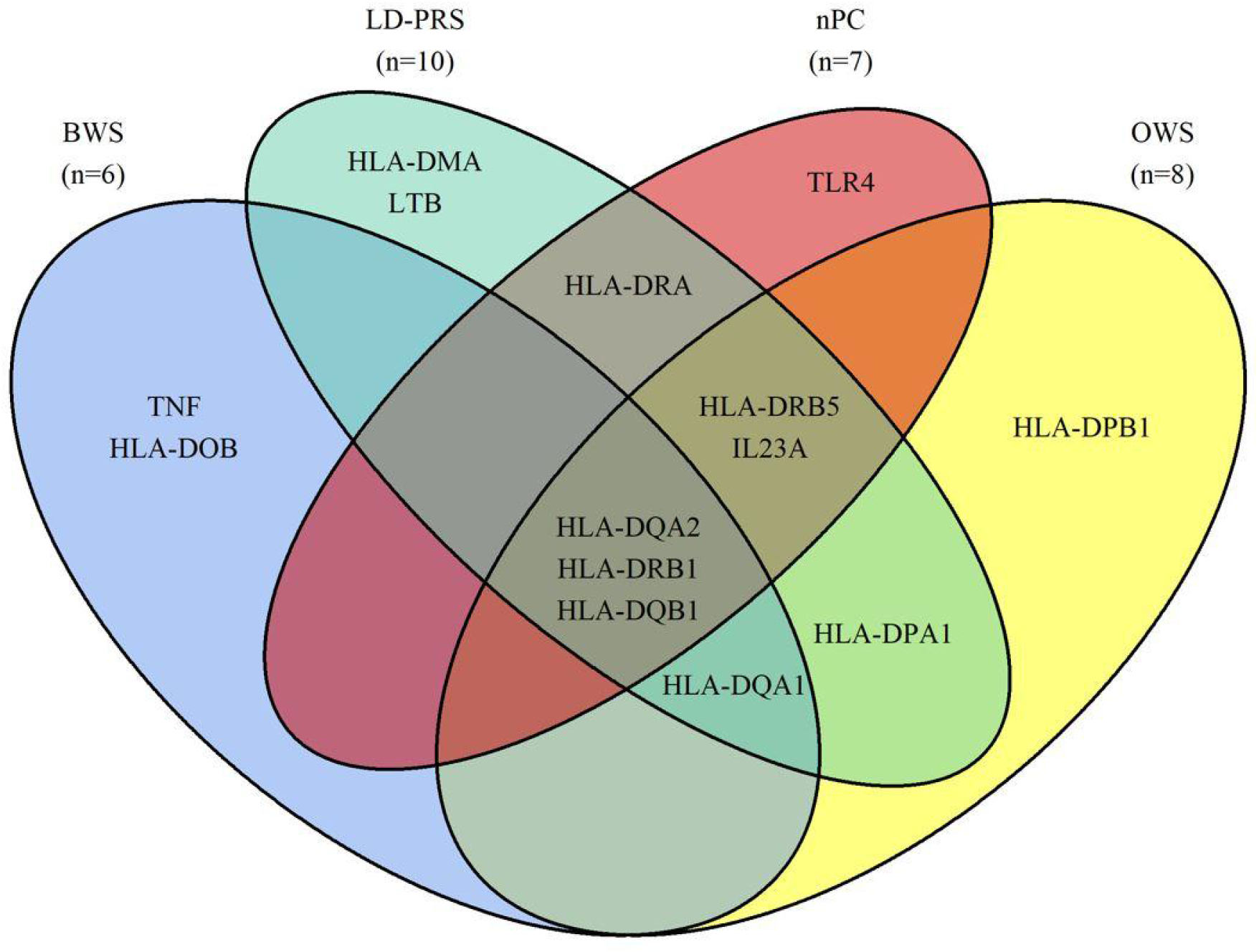
Venn diagram of the number of RA pathway genes identified by BWS, LD-PRS, OWS, and nPC for DNA methylation data.

### Application to DNA sequence data in UK Biobank

The UK Biobank is a population-based cohort study with a wide variety of genetic and phenotypic information (39). It includes ~500K individuals from the United Kingdom who are currently aged between 40 and 69 when recruited in 2006-2010 (40). We follow the same preprocess procedure in Liang et al. (41) to exclude individuals who self-report themselves not from a white British ancestry, who are marked as outliers for heterozygosity or missing rates, who have been identified to have ten or more third-degree relatives, and who are recommended for removal by the UK Biobank. Meanwhile, the quality control (QC) for DNA sequence data is also performed on both SNPs and samples using PLINK 1.9 (42) (https://www.cog-genomics.org/plink/1.9/). We filter out SNPs with missing rates larger than 5% and Hardy-Weinberg equilibrium exact test p-values less than 10^− 6^ . We also exclude individuals with missing rates larger than 5% and without sex. After QC of DNA sequence data and preprocess of phenotype data, there are 583,386 SNPs and 322,607 individuals remaining. In our analysis, we use 4,541 individuals with RA disease and randomly select 5,459 individuals without RA disease.

We define a gene to include all of the SNPs from 20 kb upstream to 20 kb downstream of the gene. We find that only 10,907 linked genes matched with genes in the above biological network that contain at least one SNP. Then, we calculate gene-level signals using OWS, LD-PRS, BWS, and nPC, respectively. The selection probability for each gene can be obtained by using half-sample method 500 times and the network-based regularization method across 600 pairs of tuning parameters *λ* and *α*. The number of genes with selection probabilities of 1 for DNA sequence data is larger than that of DNA methylation data. For example, there are 80 genes with SP=1 using OWS and 135 genes with SP=1 using LD-PRS. Therefore, we select the top 200 genes according to the selection probabilities of each method for DNA sequence data analysis. We also search the GWAS catalog (https://www.ebi.ac.uk/gwas/) for genes that are associated with RA. Fig 5 and Table 2 show the genes in the GWAS catalog that are also identified by OWS, LD-PRS, BWS, and nPC. Similar to DNA methylation data analyses, LD-PRS identifies the largest number of genes (n=23) reported in the GWAS catalog, including four uniquely identified genes (*HLA-DQB1*, *GFRA1*, *GABBR2*, *EDIL3*); OWS identify 22 genes in which genes *STAT4* (SP=0.994) and *IKZF1* (SP=0.986) are uniquely selected. There are 13 genes identified by both LD-PRS and OWS, where 12 genes have selection probabilities of 1 in both methods. Two unsupervised methods, BWS and nPC, can identify 17 and 18 genes in the GWAS catalog. They can uniquely identify 11 and 12 genes, respectively. Moreover, there are two genes identified by all four methods, genes *HLA-DQA1* and *HLA-DRA* (boldfaced in Table 2), and two genes identified by three proposed methods, genes *RATB* and *CTNNA3*.

**Fig 5.**
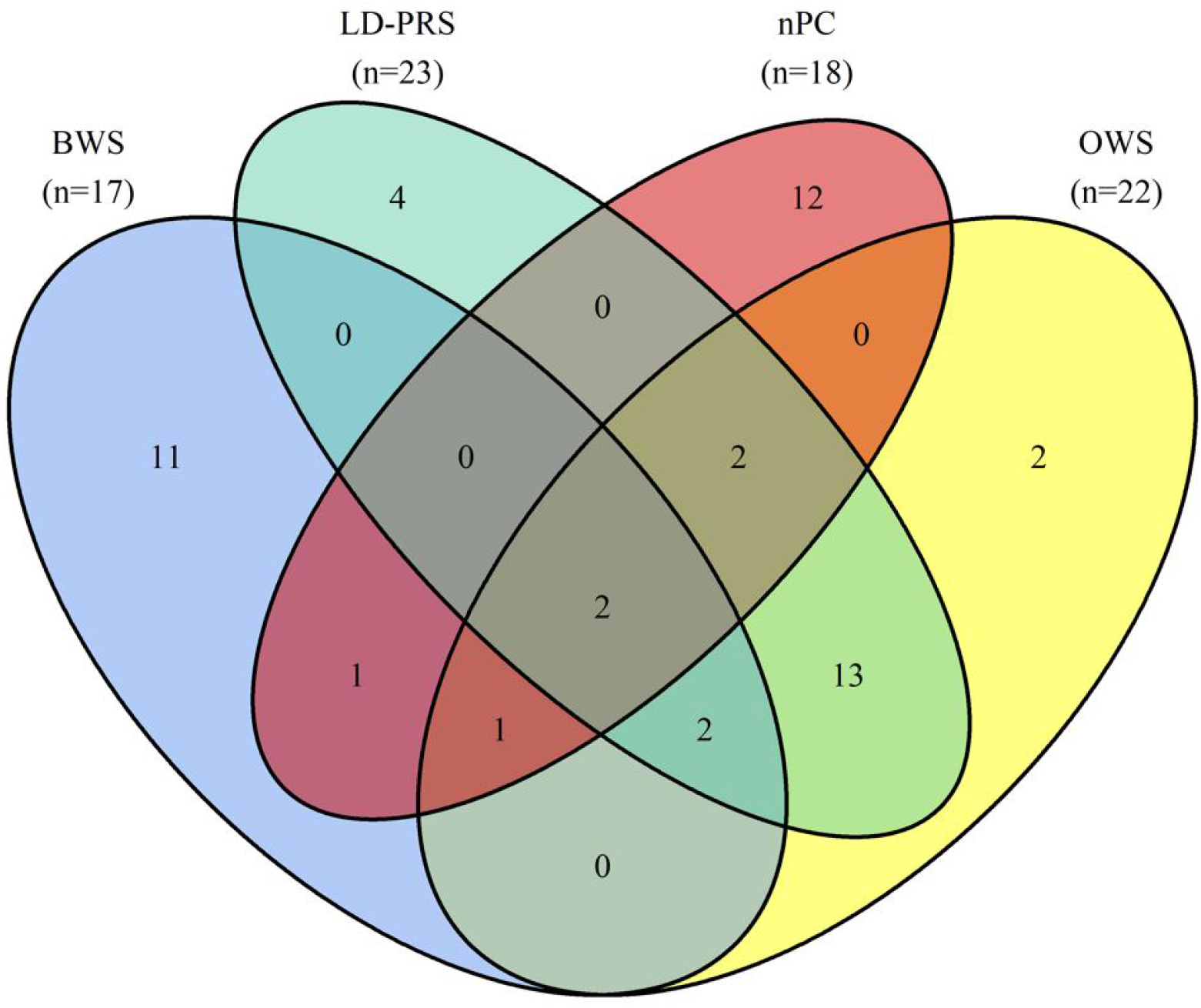
Venn diagram of the number of overlapping genes among the top 200 genes identified by each method and reported in the GWAS catalog for DNA sequence data.

**Table 2.**
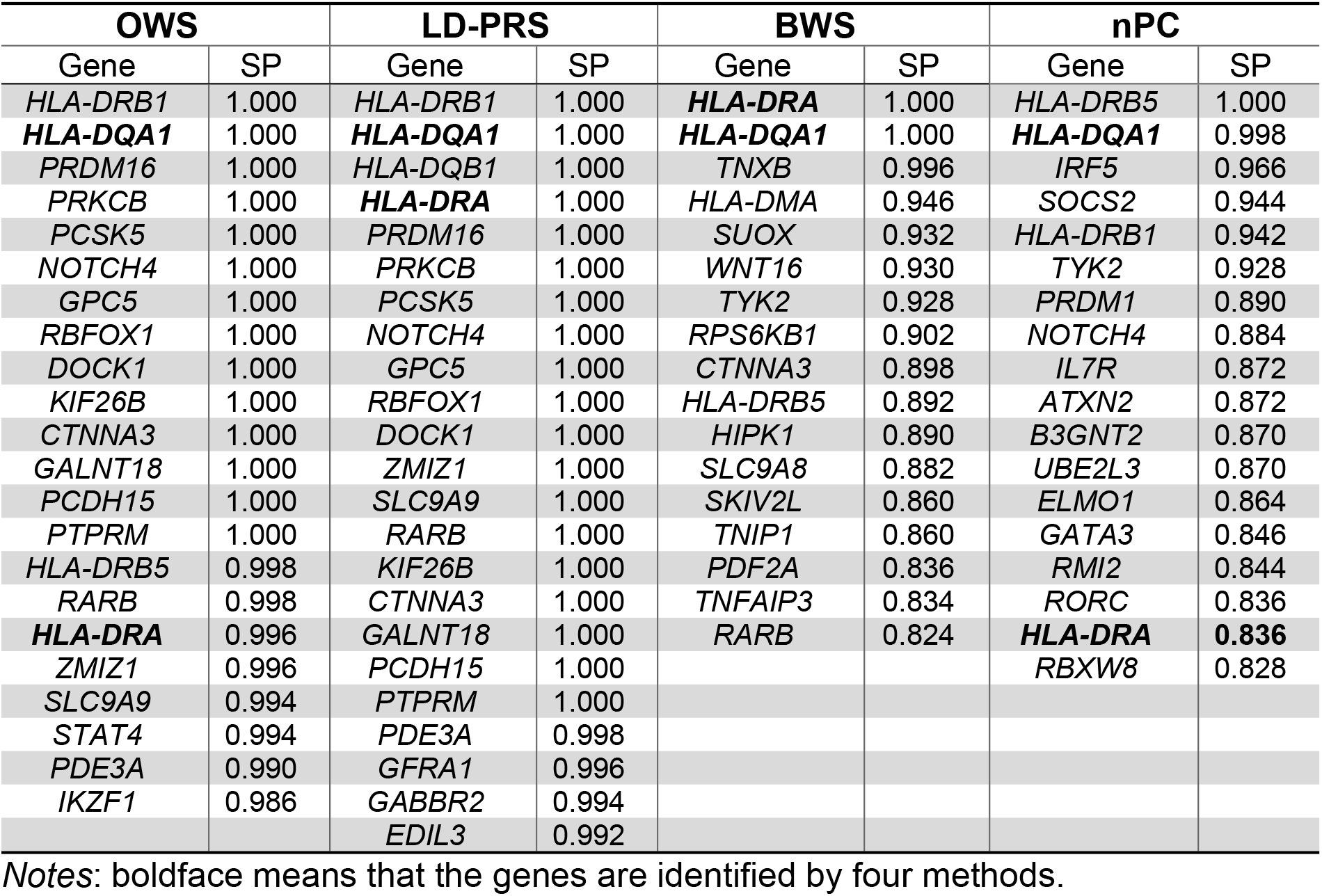
GWAS catalog reported genes identified by OWS, LD-PRS, BWS, and nPC in DNA sequence data.

## Discussions

In this paper, we develop three linear combination approaches to capture the gene-level signals from multiple CpG sites or SNPs: optimally weighted sum (OWS), LD-adjusted polygenic risk score (LD-PRS), and beta-based weighted sum (BWS) in DNA methylation or DNA sequence data. To identify phenotype related genes, we apply the three gene-level signals to a stability gene selection approach by incorporating genetic networks. Compared with the traditional dimension reduction techniques such as PC based gene-level signal, our proposed approaches have very good performance according to the true positive rates. By applying our methods to real DNA methylation and DNA sequence data, we show that the newly proposed approaches can select more potentially RA related genes that are missed by nPC. Meanwhile, our proposed methods can select more significantly enriched genes in the RA pathway comparing with nPC, such as genes *HLA-DMA*, *HLA-DPB1*, and *HLA-DOB* in the HLA region.

There are some advantages of our proposed gene-level signals. First, the three linear combination approaches can capture more information from genetic components (SNPs or CpG sites) in a gene than the traditional dimension reduction techniques, such as PC-based methods. OWS and LD-PRS are two supervised approaches based on the association between each genetic component and the phenotype, where OWS utilizes the optimally weighted combination (12) of components and LD-PRS can adjust for the highly correlated structure (13) of components. OWS puts large weights on components with large effects on the phenotype (12). Since the genetic components in a gene are commonly correlated, LD-PRS transforms the original data into an orthogonal space to adjust for LD structure. Moreover, OWS and LD-PRS perform better according to TPR when the genetic components are highly correlated. Even though BWS is an unsupervised method that can be extracted without using phenotype, our simulation studies show that BWS has the highest TPR in most of the settings. Second, the proposed methods can select more potential phenotype related genes. In our application to DNA methylation of RA patients and normal controls, the top 100 genes selected by our proposed methods can be significantly enriched into RA pathway and contain more RA pathway genes, especially by LD-PRS. Furthermore, all of our proposed methods have strong evidence to select gene *HLA-QDA2* (SP=1) which is not reported in the GWAS catalog.

Recently, large-scale biobanks linked to electronic health records provide us the possibility of analyzing DNA sequence data using a large sample size. Although our proposed linear combination approaches combined with the network regularization have several advantages, there are three limitations we need to resolve in our future works. First, the proposed methods are not suitable for extremely unbalanced case-control studies. To avoid the extremely unbalanced case-control ratio in the data from UK Biobank, we match the number of individuals with and without RA disease in the application of DNA sequence data. This may be the reason for a large number of genes with SP=1 using OWS and LD-PRS, and the SP of the 200^th^ gene using OWS and LD-PRS over 0.97. The second limitation is that we do not know if the genes selected by the proposed methods are significantly associated with the phenotype. For future studies, we plan to integrate statistical inference in the selection procedure, and further investigate the selection performance by integrating both selection and statistical inference. The third limitation is that the network-based regularization method is only used for case-control study (9). For the continuous phenotypes, we need to switch the logistic model with logistic likelihood to the linear regression model with mean squared error or more robust loss function, such as Huber function (43).

## AUTHOR CONTRIBUTION

Formal analysis: Xuewei Cao; Methodology: Xuewei Cao, Shanglin Zhang, Xiaoyu Liang, and Qiuying Sha; Data curation: Xuewei Cao and Xiaoyu Liang; Visualization: Xuewei Cao; Writing original draft: Xuewei Cao, Xiaoyu Liang, and Qiuying Sha; Writing review and editing: Xuewei Cao, Shuanglin Zhang, Xiaoyu Liang, and Qiuying Sha.

## ACKNOWLEDGEMENTS

Part of this research has been conducted using the UK Biobank Resource under application number 41722 and the NHGRI-EBI GWAS Catalog.

## Supporting information

**S1 Fig.**
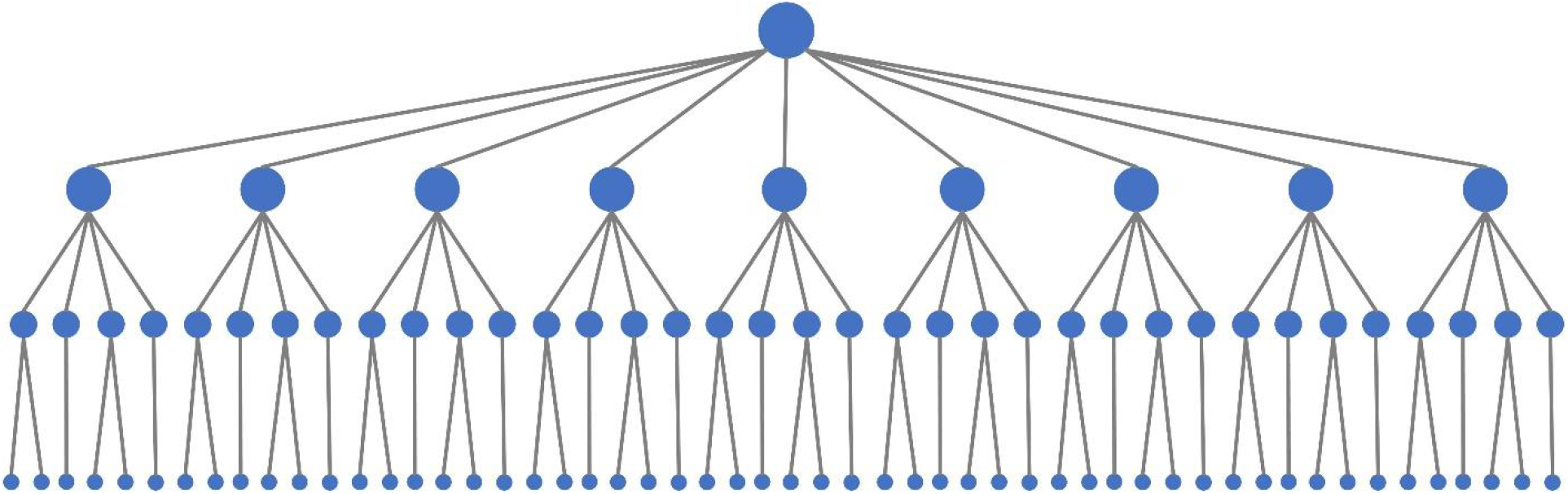
The network module used in simulation studies. There are a total of 100 genes which contain a centered gene.

**S2 Fig.**
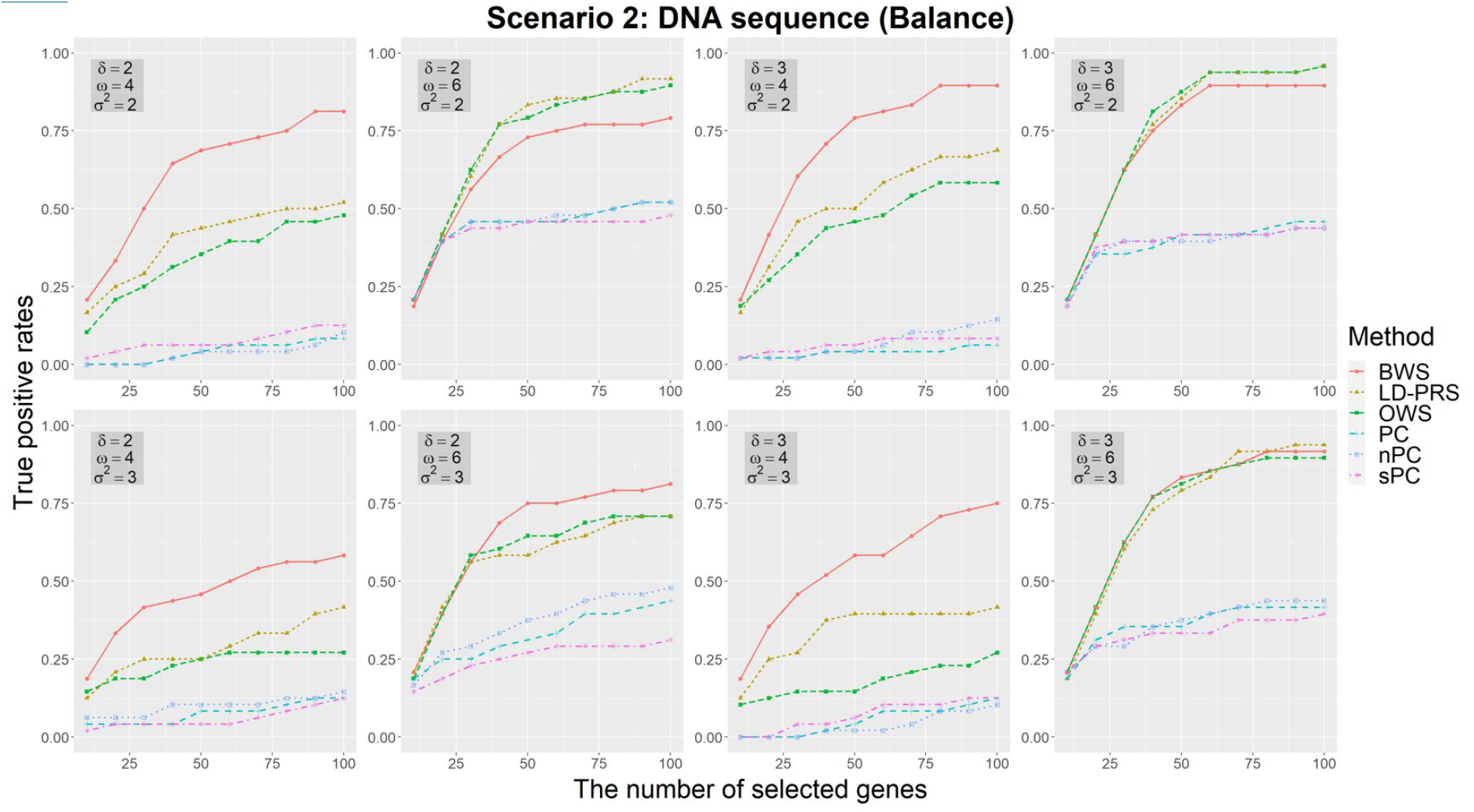
The true positive rates of the methods based on different gene-level signals for balance case-control studies with DNA sequence data in scenario 2, where there are five rare variants and five common variants in each gene. According to the different number of selected top-genes, three parameters are used to vary the genetic effect: the strength of association signals *δ*, the number of SNPs in each gene related to gene-level signals *ω*, and the noise level of association signals *σ*^2^. The selection probabilities are calculated using half-sample method 100 times.

**S3 Fig.**
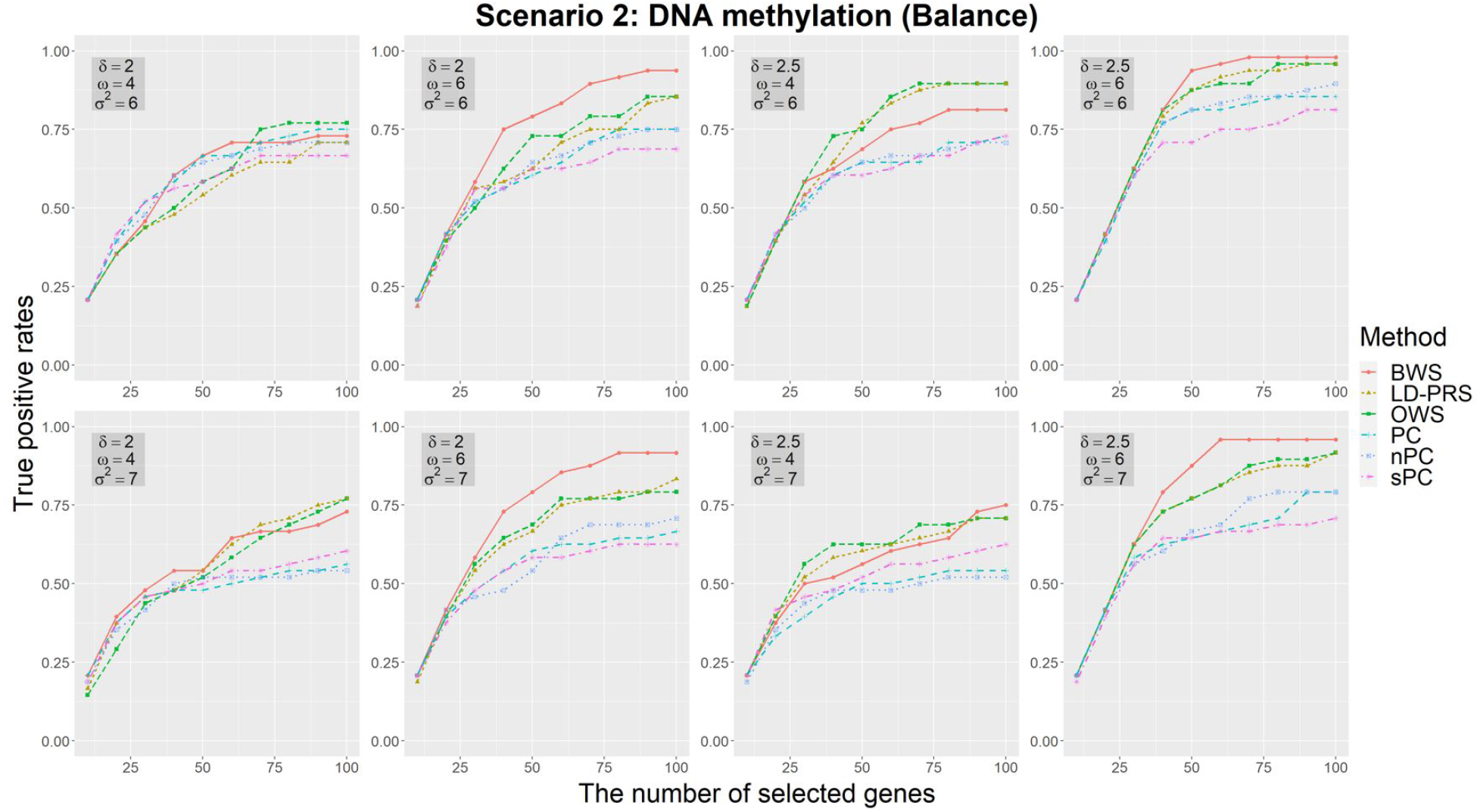
The true positive rates of the methods based on different gene-level signals for balance case-control studies with DNA methylation data in scenario 2. According to the different number of selected top-genes, three parameters are used to vary the genetic effect: the strength of association signals *δ*, the number of CpG sites in each gene related to gene-level signals *ω*, and the noise level of association signals *σ*^2^. The selection probabilities are calculated using half-sample method 100 times.

**S4 Fig.**
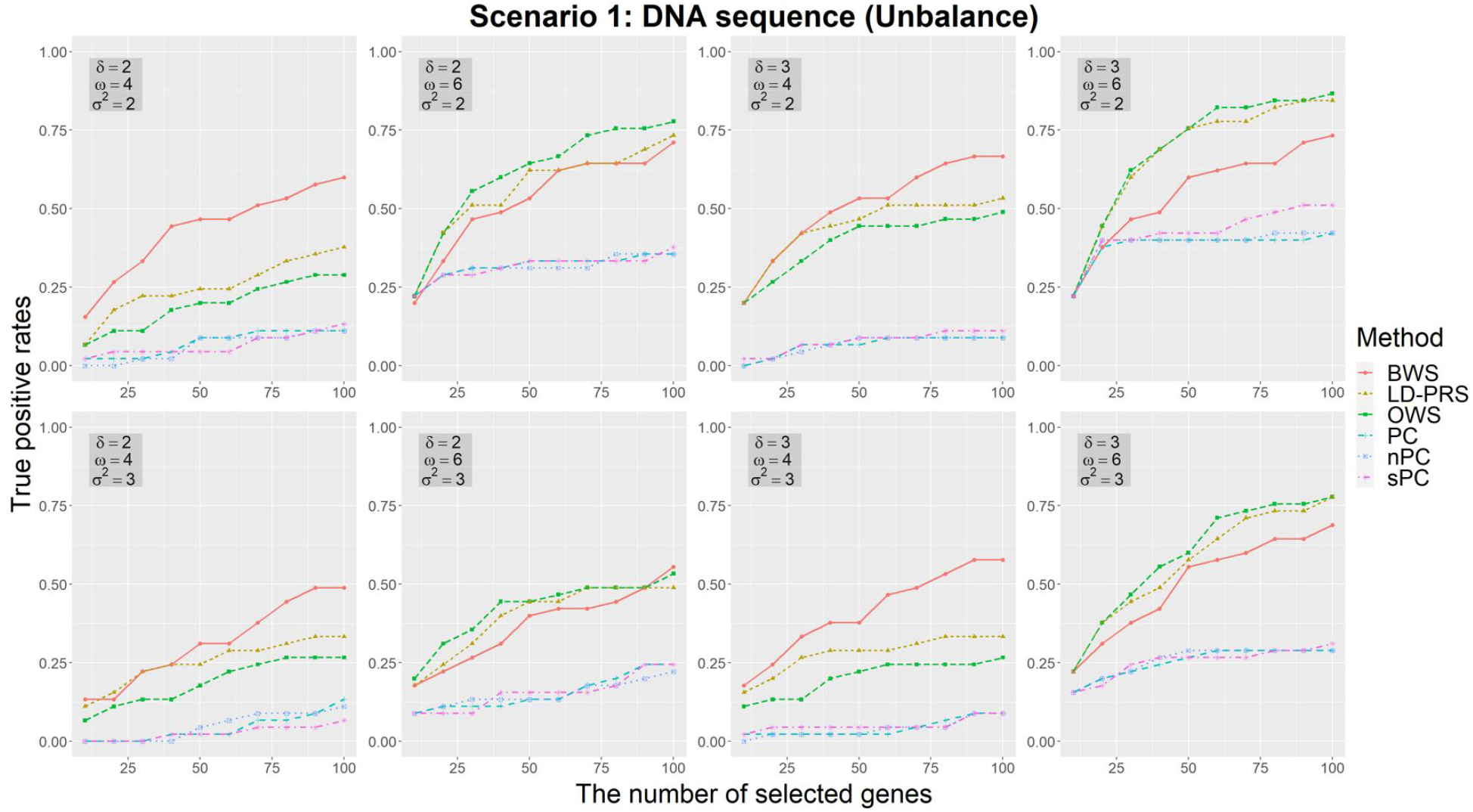
The true positive rates of the methods based on different gene-level signals for unbalance case-control studies (case:control=100:900) with DNA sequence data in scenario 1, where there are five rare variants and five common variants in each gene. According to the different number of selected top-genes, three parameters are used to vary the genetic effect: the strength of association signals *δ*, the number of SNPs in each gene related to gene-level signals *ω*, and the noise level of association signals *σ*^2^. The selection probabilities are calculated using half-sample method 100 times.

**S5 Fig.**
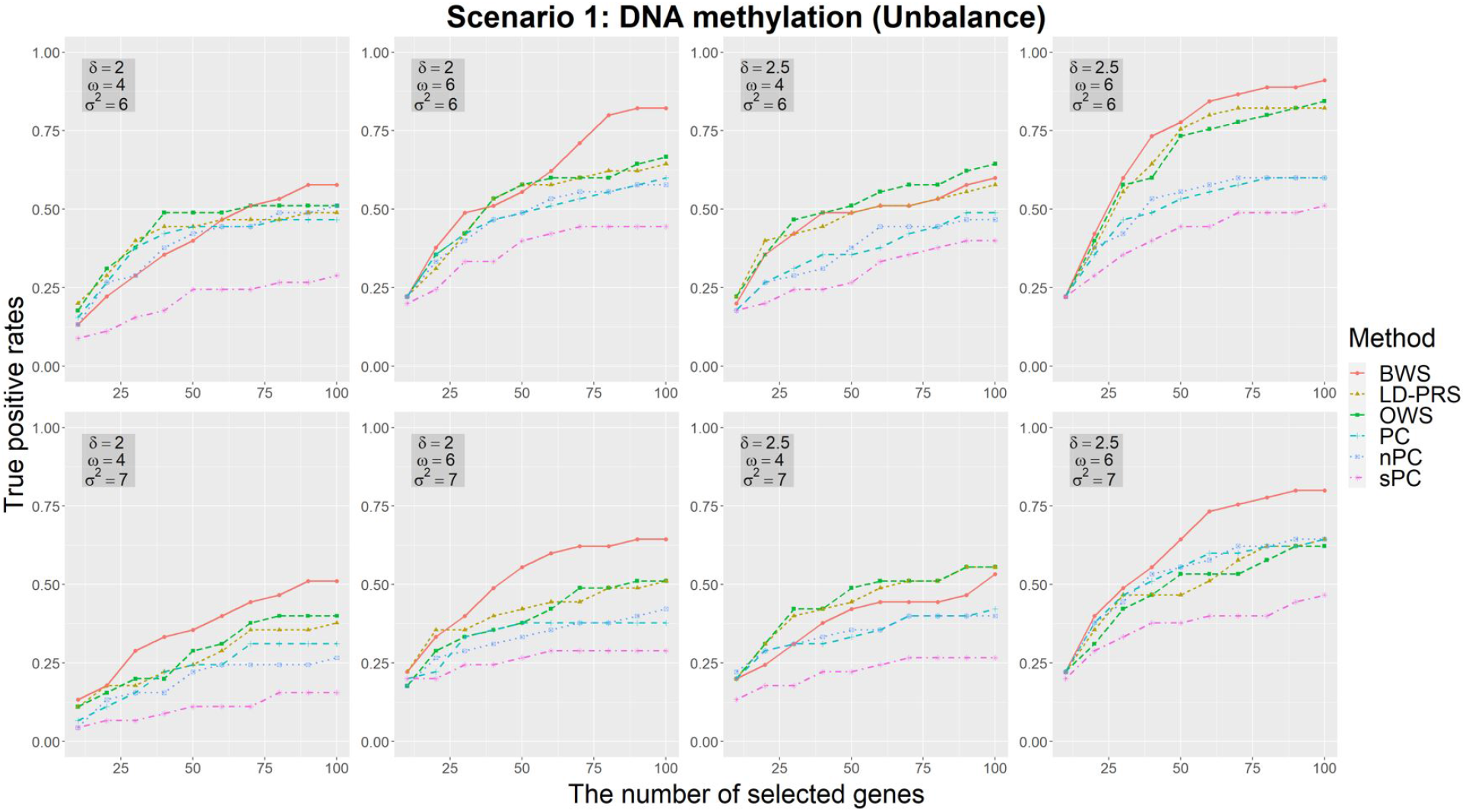
The true positive rates of the methods based on different gene-level signals for unbalance case-control studies (case:control=100:900) with DNA methylation data in scenario 1. According to the different number of selected top-genes, three parameters are used to vary the genetic effect: the strength of association signals *δ*, the number of CpG sites in each gene related to gene-level signals *ω*, and the noise level of association signals *σ*^2^. The selection probabilities are calculated using half-sample method 100 times.

**S6 Fig.**
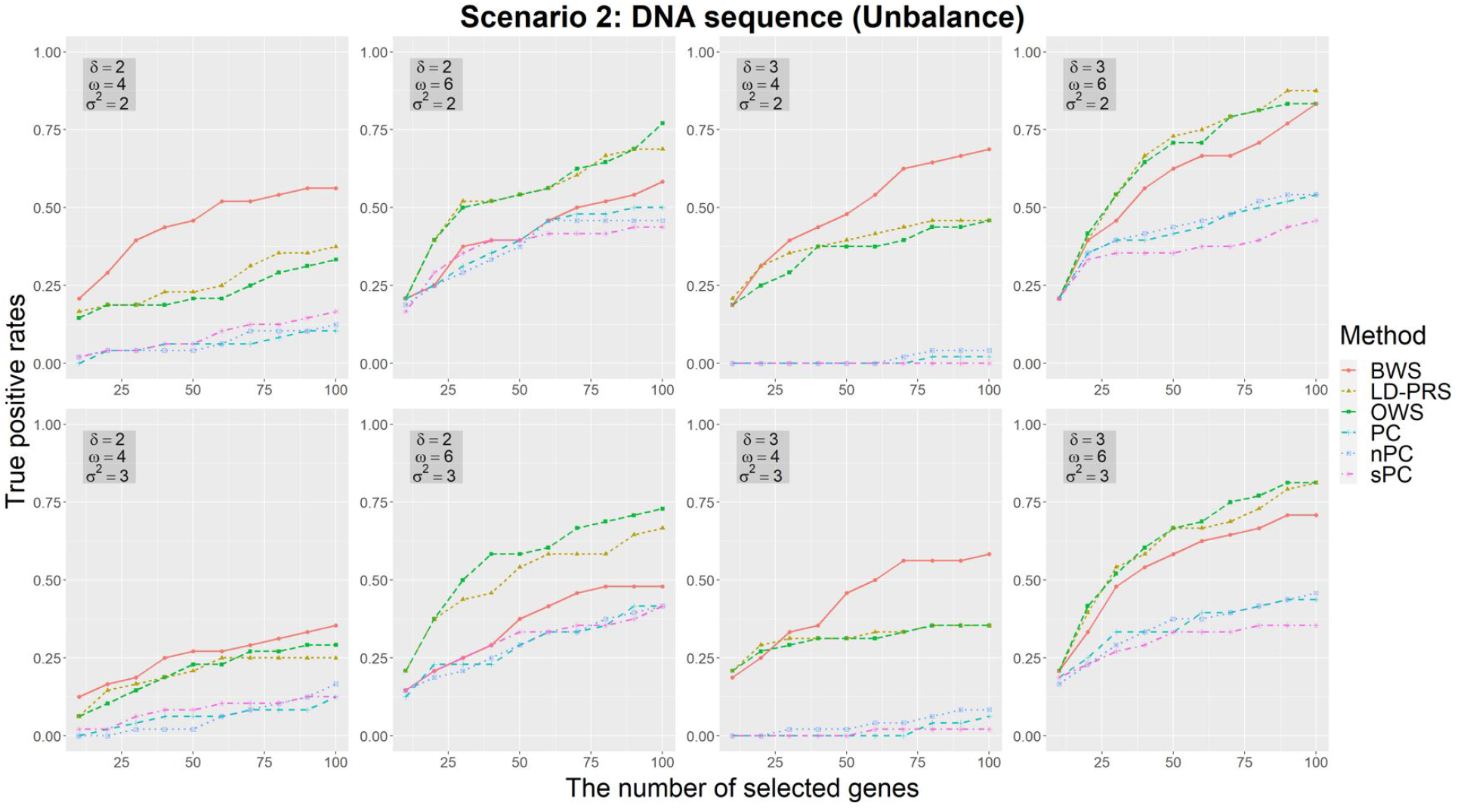
The true positive rates of the methods based on different gene-level signals for unbalance case-control studies (case:control=100:900) with DNA sequence data in scenario 2, where there are five rare variants and five common variants in each gene. According to the different number of selected top-genes, three parameters are used to vary the genetic effect: the strength of association signals *δ*, the number of SNPs in each gene related to gene-level signals *ω*, and the noise level of association signals *σ*^2^. The selection probabilities are calculated using half-sample method 100 times.

**S7 Fig.**
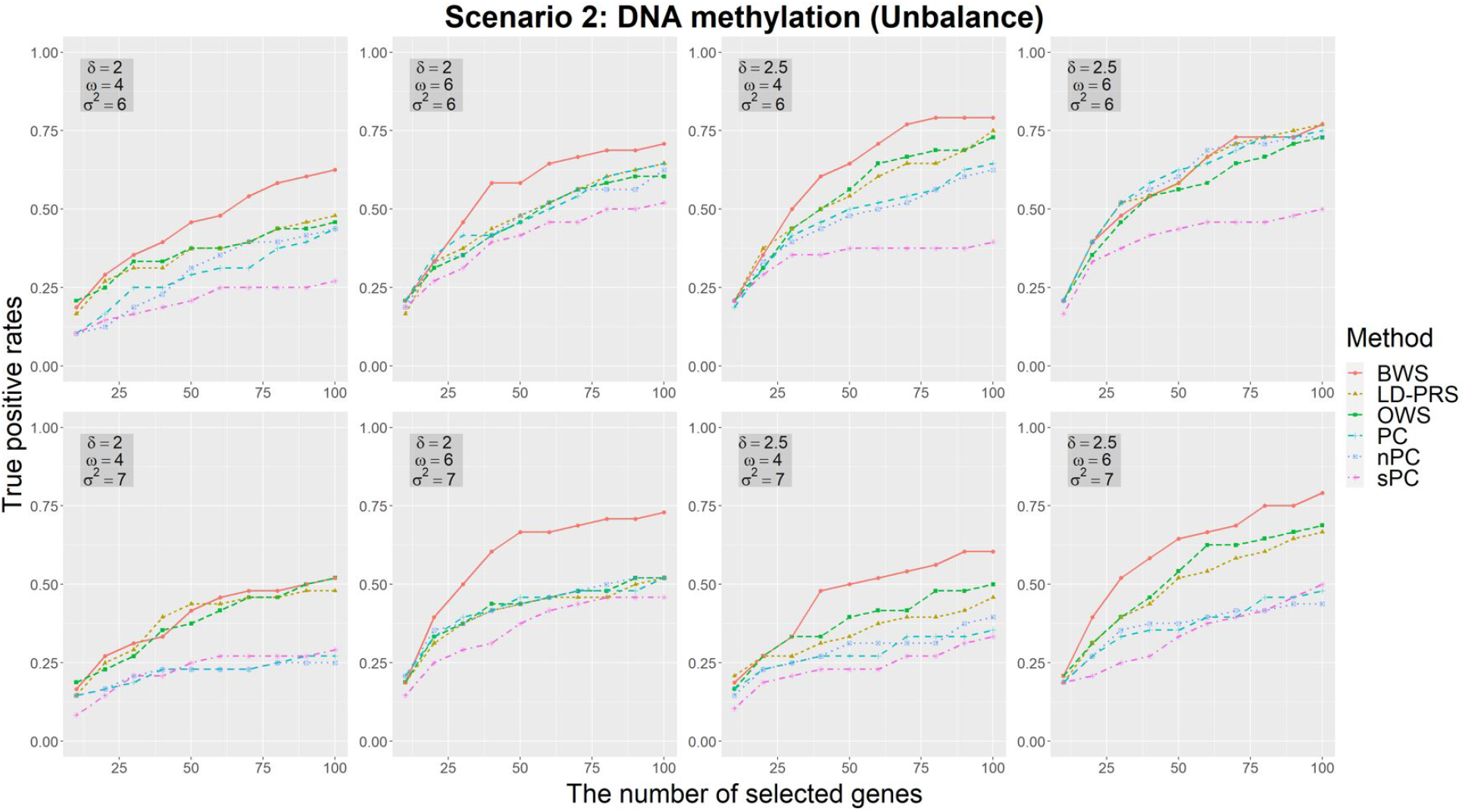
The true positive rates of the methods based on different gene-level signals for unbalance case-control studies (case:control=100:900) with DNA methylation data in scenario 2. According to the different number of selected top-genes, three parameters are used to vary the genetic effect: the strength of association signals *δ*, the number of CpG sites in each gene related to gene-level signals *ω*, and the noise level of association signals *σ*^2^. The selection probabilities are calculated using half-sample method 100 times.

**S8 Fig.**
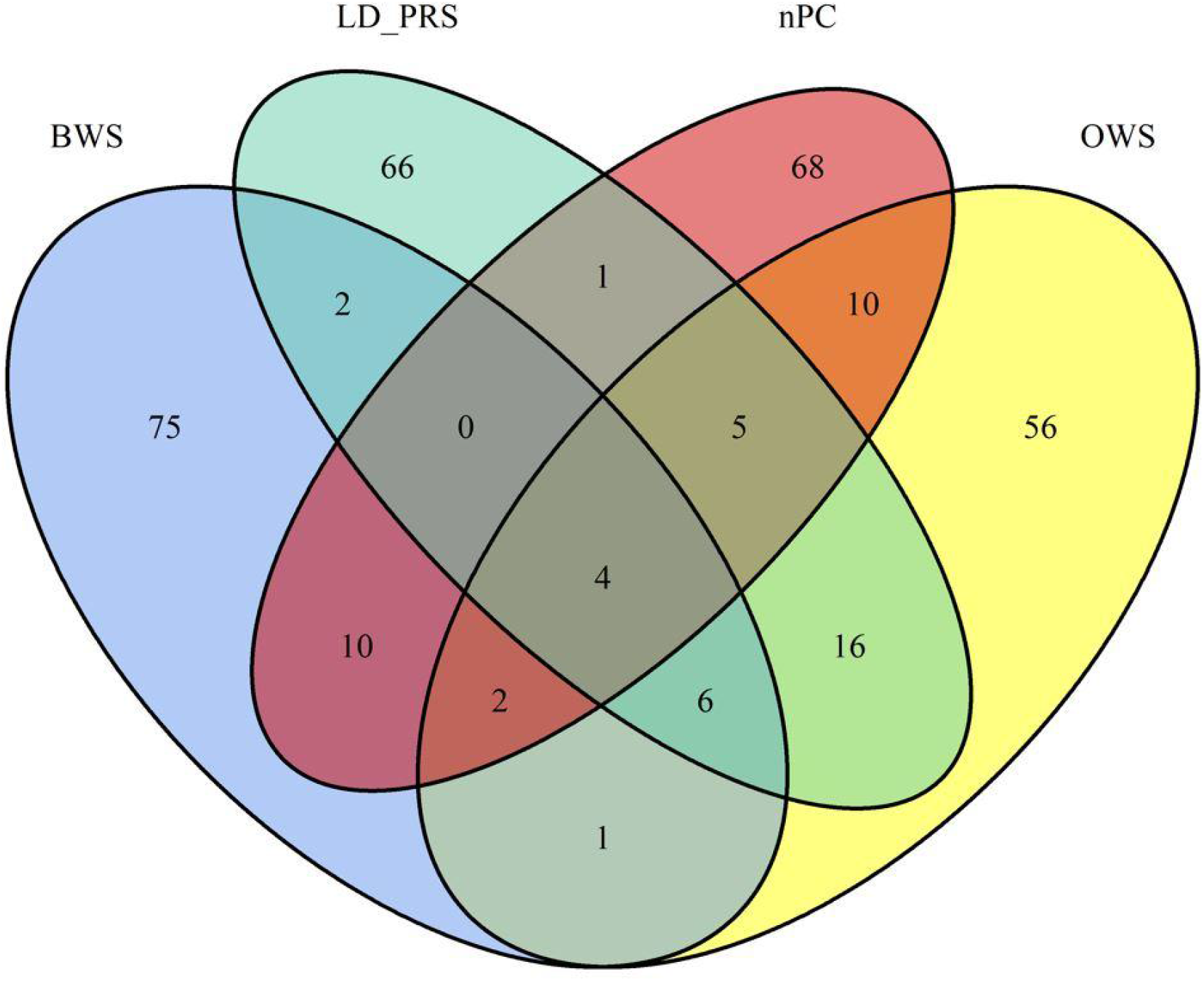
Venn diagram of the number of top 100 genes identified by BWS, LD-PRS, OWS, and nPC for DNA methylation data.

## References

1. Ritchie MD. Large-scale analysis of genetic and clinical patient data. Annual Review of Biomedical Data Science. 2018;1:263–74.

2. Li R, Duan R, Kember RL, Rader DJ, Damrauer SM, Moore JH, et al. A regression framework to uncover pleiotropy in large-scale electronic health record data. Journal of the American Medical Informatics Association. 2019;26(10):1083–90.

3. Wang DG, Fan J-B, Siao C-J, Berno A, Young P, Sapolsky R, et al. Large-scale identification, mapping, and genotyping of single-nucleotide polymorphisms in the human genome. Science. 1998;280(5366):1077–82.

4. Bock C. Analysing and interpreting DNA methylation data. Nature Reviews Genetics. 2012;13(10):705–19.

5. Waldmann P, Mészáros G, Gredler B, Fuerst C, Sölkner J. Evaluation of the lasso and the elastic net in genome-wide association studies. Frontiers in genetics. 2013;4:270.

6. Wang H, Lengerich BJ, Aragam B, Xing EP. Precision Lasso: accounting for correlations and linear dependencies in high-dimensional genomic data. Bioinformatics. 2019;35(7):1181–7.

7. Yuan M, Lin Y. Model selection and estimation in regression with grouped variables. Journal of the Royal Statistical Society: Series B (Statistical Methodology). 2006;68(1):49–67.

8. Meier L, Van De Geer S, Bühlmann P. The group lasso for logistic regression. Journal of the Royal Statistical Society: Series B (Statistical Methodology). 2008;70(1):53–71.

9. Kim K, Sun H. Incorporating genetic networks into case-control association studies with high-dimensional DNA methylation data. BMC bioinformatics. 2019;20(1):1–15.

10. Li C, Li H. Network-constrained regularization and variable selection for analysis of genomic data. Bioinformatics. 2008;24(9):1175–82.

11. Sun H, Wang S. Network-based regularization for matched case-control analysis of high-dimensional DNA methylation data. Statistics in medicine. 2013;32(12):2127–39.

12. Sha Q, Wang X, Wang X, Zhang S. Detecting association of rare and common variants by testing an optimally weighted combination of variants. Genetic epidemiology. 2012;36(6):561–71.

13. Baker E, Schmidt KM, Sims R, O'Donovan MC, Williams J, Holmans P, et al. POLARIS: Polygenic LD-adjusted risk score approach for set-based analysis of GWAS data. Genetic epidemiology. 2018;42(4):366–77.

14. Wu MC, Lee S, Cai T, Li Y, Boehnke M, Lin X. Rare-variant association testing for sequencing data with the sequence kernel association test. The American Journal of Human Genetics. 2011;89(1):82–93.

15. Li C, Li H. Variable selection and regression analysis for graph-structured covariates with an application to genomics. The annals of applied statistics. 2010;4(3):1498.

16. Choi J, Kim K, Sun H. New variable selection strategy for analysis of high-dimensional DNA methylation data. Journal of bioinformatics and computational biology. 2018;16(04):1850010.

17. Sun H, Wang S. Penalized logistic regression for high-dimensional DNA methylation data with case-control studies. Bioinformatics. 2012;28(10):1368–75.

18. Meinshausen N, Bühlmann P. Stability selection. Journal of the Royal Statistical Society: Series B (Statistical Methodology). 2010;72(4):417–73.

19. Friedman J, Hastie T, Tibshirani R. Regularization paths for generalized linear models via coordinate descent. Journal of statistical software. 2010;33(1):1.

20. Peng J, Wang P, Zhou N, Zhu J. Partial correlation estimation by joint sparse regression models. Journal of the American Statistical Association. 2009;104(486):735–46.

21. Liu Y, Aryee MJ, Padyukov L, Fallin MD, Hesselberg E, Runarsson A, et al. Epigenome-wide association data implicate DNA methylation as an intermediary of genetic risk in rheumatoid arthritis. Nature biotechnology. 2013;31(2):142–7.

22. Kular L, Liu Y, Ruhrmann S, Zheleznyakova G, Marabita F, Gomez-Cabrero D, et al. DNA methylation as a mediator of HLA-DRB1* 15: 01 and a protective variant in multiple sclerosis. Nature communications. 2018;9(1):1–15.

23. Jiang X, Källberg H, Chen Z, Ärlestig L, Rantapää-Dahlqvist S, Davila S, et al. An Immunochip-based interaction study of contrasting interaction effects with smoking in ACPA-positive versus ACPA-negative rheumatoid arthritis. Rheumatology. 2016;55(1):149–55.

24. Traylor M, Knevel R, Cui J, Taylor J, Harm-Jan W, Conaghan PG, et al. Genetic associations with radiological damage in rheumatoid arthritis: Meta-analysis of seven genome-wide association studies of 2,775 cases. PloS one. 2019;14(10):e0223246.

25. Eyre S, Bowes J, Diogo D, Lee A, Barton A, Martin P, et al. High-density genetic mapping identifies new susceptibility loci for rheumatoid arthritis. Nature genetics. 2012;44(12):1336–40.

26. Govind N, Choudhury A, Hodkinson B, Ickinger C, Frost J, Lee A, et al. Immunochip identifies novel, and replicates known, genetic risk loci for rheumatoid arthritis in black South Africans. Molecular Medicine. 2014;20(1):341–9.

27. Plenge RM, Seielstad M, Padyukov L, Lee AT, Remmers EF, Ding B, et al. TRAF1–C5 as a risk locus for rheumatoid arthritis—a genomewide study. New England Journal of Medicine. 2007;357(12):1199–209.

28. Bossini-Castillo L, De Kovel C, Kallberg H, van‘t Slot R, Italiaander A, Coenen M, et al. A genome-wide association study of rheumatoid arthritis without antibodies against citrullinated peptides. Annals of the rheumatic diseases. 2015;74(3):e15–e.

29. Consortium WTCC. Genome-wide association study of 14,000 cases of seven common diseases and 3,000 shared controls. Nature. 2007;447(7145):661.

30. Wei W-H, Viatte S, Merriman TR, Barton A, Worthington J. Genotypic variability based association identifies novel non-additive loci DHCR7 and IRF4 in sero-negative rheumatoid arthritis. Scientific reports. 2017;7(1):1–7.

31. Julia A, Ballina J, Canete JD, Balsa A, Tornero-Molina J, Naranjo A, et al. Genome-wide association study of rheumatoid arthritis in the Spanish population: KLF12 as a risk locus for rheumatoid arthritis susceptibility. Arthritis & Rheumatism: Official Journal of the American College of Rheumatology. 2008;58(8):2275–86.

32. Negi S, Juyal G, Senapati S, Prasad P, Gupta A, Singh S, et al. A genome-wide association study reveals ARL15, a novel non-HLA susceptibility gene for rheumatoid arthritis in North Indians. Arthritis & Rheumatism. 2013;65(12):3026–35.

33. Aterido A, Cañete JD, Tornero J, Ferrándiz C, Pinto JA, Gratacós J, et al. Genetic variation at the glycosaminoglycan metabolism pathway contributes to the risk of psoriatic arthritis but not psoriasis. Annals of the Rheumatic diseases. 2019;78(3):355–64.

34. Kochi Y, Okada Y, Suzuki A, Ikari K, Terao C, Takahashi A, et al. A regulatory variant in CCR6 is associated with rheumatoid arthritis susceptibility. Nature genetics. 2010;42(6):515–9.

35. Raychaudhuri S, Remmers EF, Lee AT, Hackett R, Guiducci C, Burtt NP, et al. Common variants at CD40 and other loci confer risk of rheumatoid arthritis. Nature genetics. 2008;40(10):1216–23.

36. Weyand CM, Goronzy JJ. Association of MHC and rheumatoid arthritis: HLA polymorphisms in phenotypic variants of rheumatoid arthritis. Arthritis Research & Therapy. 2000;2(3):1–5.

37. Huang DW, Sherman BT, Lempicki RA. Bioinformatics enrichment tools: paths toward the comprehensive functional analysis of large gene lists. Nucleic acids research. 2009;37(1):1–13.

38. Sherman BT, Lempicki RA. Systematic and integrative analysis of large gene lists using DAVID bioinformatics resources. Nature protocols. 2009;4(1):44–57.

39. Bycroft C, Freeman C, Petkova D, Band G, Elliott LT, Sharp K, et al. The UK Biobank resource with deep phenotyping and genomic data. Nature. 2018;562(7726):203–9.

40. Sudlow C, Gallacher J, Allen N, Beral V, Burton P, Danesh J, et al. UK biobank: an open access resource for identifying the causes of a wide range of complex diseases of middle and old age. PLoS medicine. 2015;12(3):e1001779.

41. Liang X, Cao X, Sha Q, Zhang S. HCLC-FC: a Novel Statistical Method for Phenome-Wide Association Studies. Submitted to Bioinformatics. 2022.

42. Chang CC, Chow CC, Tellier LC, Vattikuti S, Purcell SM, Lee JJ. Second-generation PLINK: rising to the challenge of larger and richer datasets. Gigascience. 2015;4(1):s13742-015-0047-8.

43. Huber PJ. Robust estimation of a location parameter. Breakthroughs in statistics: Springer; 1992. p. 492–518.

